# Beta cell extracellular vesicle PD-L1 as a novel regulator of CD8+ T cell activity and biomarker during the evolution of Type 1 Diabetes

**DOI:** 10.1101/2024.09.18.613649

**Authors:** Chaitra Rao, Daniel T Cater, Saptarshi Roy, Jerry Xu, Andre De G Olivera, Carmella Evans-Molina, Jon D. Piganelli, Decio L. Eizirik, Raghavendra G. Mirmira, Emily K. Sims

**Author notes:** Authors contributed equally to this work (Chaitra Rao, Daniel T Cater). **Corresponding author** Emily K. Sims M.D. M.S Associate Professor of Pediatrics, Indiana University School of Medicine Center for Diabetes and Metabolic Diseases 635 Barnhill Drive, MS 2031 Indianapolis, IN 46202.

## Abstract

**Aims/hypothesis:** Surviving beta cells in type 1 diabetes respond to inflammation by upregulating programmed death-ligand 1 (PD-L1) to engage immune cell programmed death-1 (PD-1) and limit destruction by self-reactive immune cells. Extracellular vesicles (EVs) and their cargo can serve as biomarkers of beta cell health and contribute to islet intercellular communication. We hypothesized that the inflammatory milieu of type 1 diabetes increases PD-L1 in beta cell EV cargo and that EV PD-L1 may protect beta cells against immune-mediated cell death.

**Methods:** Beta cell lines and human islets were treated with proinflammatory cytokines to model the proinflammatory type 1 diabetes microenvironment. EVs were isolated using ultracentrifugation or size exclusion chromatography and analysed via immunoblot, flow cytometry, and ELISA. EV PD-L1: PD-1 binding was assessed using a competitive binding assay and *in vitro* functional assays testing the ability of EV PD-L1 to inhibit NOD CD8 T cells. Plasma EV and soluble PD-L1 were assayed in plasma of individuals with islet autoantibody positivity (Ab+) or recent-onset type 1 diabetes and compared to non-diabetic controls.

**Results:** PD-L1 protein colocalized with tetraspanin-associated proteins intracellularly and was detected on the surface of beta cell EVs. 24-h IFN-α or IFN-□ treatment induced a two-fold increase in EV PD-L1 cargo without a corresponding increase in number of EVs. IFN exposure predominantly increased PD-L1 expression on the surface of beta cell EVs and beta cell EV PD-L1 showed a dose-dependent capacity to bind PD-1. Functional experiments demonstrated specific effects of beta cell EV PD-L1 to suppress proliferation and cytotoxicity of murine CD8 T cells. Plasma EV PD-L1 levels were increased in islet Ab+ individuals, particularly in those with single Ab+, Additionally, in from individuals with either Ab+ or type 1 diabetes, but not in controls, plasma EV PD-L1 positively correlated with circulating C-peptide, suggesting that higher EV-PD-L1 could be protective for residual beta cell function.

**Conclusions/interpretation:** IFN exposure increases PD-L1 on the beta cell EV surface. Beta cell EV PD-L1 binds PD1 and inhibits CD8 T cell proliferation and cytotoxicity. Circulating EV PD-L1 is higher in islet autoantibody positive patients compared to controls. Circulating EV PD-L1 levels correlate with residual C-peptide at different stages in type 1 diabetes progression. These findings suggest that EV PD-L1 could contribute to heterogeneity in type 1 diabetes progression and residual beta cell function and raise the possibility that EV PD-L1 could be exploited as a means to inhibit immune-mediated beta cell death.

**Research in context:** *What is already known about this subject? (maximum of 3 bullet points):*

- Extracellular vesicles (EVs) serve as paracrine effectors in the islet microenvironment in health and disease.
- Interferon-alpha (IFN-α) and IFN-gamma (IFN-□) are key cytokines contributing to type 1 diabetes pathophysiology and islet IFN signalling increases beta cell programmed death-ligand 1 (PD-L1) expression.
- Up-regulation of beta cell PD-L1 in the non-obese diabetic (NOD) mouse model delays progression of type 1 diabetes.

*What is the key question? (one bullet point only; formatted as a question):*

- Do beta cells exposed to IFNs upregulate EV PD-L1 and can these changes be detected in circulation?

*What are the new findings? (maximum of 3 bullet points):*

- IFN-α or IFN-□ exposure increases beta cell EV PD-L1 cargo in beta cell lines and human islets.
- PD-L1 is present on the surface of beta cell EVs, binds PD-1 and EV PD-L1 inhibits proliferation, activation and cytotoxicity of murine CD8 T cells.
- EV PD-L1 levels are higher in islet autoantibody positive individuals compared to nondiabetic controls and levels of circulating EV PD-L1 positively correlate with residual beta cell function in islet autoantibody positive individuals as well as in individuals with recent-onset type 1 diabetes.

*How might this impact on clinical practice in the foreseeable future? (one bullet point only):*

- A beneficial effect of PD-L1+ EVs could ultimately be harnessed as an intervention to prevent autoimmune beta cell destruction. Circulating EV PD-L1 cargo has potential as a minimally invasive and informative biomarker to offer insights into the pathogenesis and progression of type 1 diabetes.

## Introduction

The global prevalence of type 1 diabetes is increasing, with current modelling predicting a prevalence of 13.5-17.4 million individuals by 2040 [1–3]. Mechanistic understanding of factors impacting disease natural history is critical to understanding heterogeneity in type 1 diabetes progression as well as the development of disease-modifying therapies. An increasing body of evidence demonstrates a critical role of intrinsic beta cell signalling in the potentiation of immune cell pathogenicity in type 1 diabetes [4–6]. One such pathway involves immune checkpoints, such as programmed death-ligand 1 (PD-L1), a normal component of the immune system that provides negative feedback on immune cells [7]. Here, PD-L1, found on the surface of cells potentially targeted by the immune system, binds its receptor programmed cell death protein 1 (PD-1) on immune cells to inhibit their activity [8]. Immune checkpoint inhibitors, including PD-L1 and PD-1 inhibitors, are used in certain cancers to increase antitumor immune responses against cancer cells [9]. Beta cells also express PD-L1, and intriguingly, while checkpoint inhibitor treatment frequently results in autoimmune complications including type 1 diabetes, PD-L1 up-regulation prevents diabetes in mouse models [10–14]. These observations suggest that PD-L1 signalling plays a critical role in beta cell survival in type 1 diabetes.

Extracellular vesicles (EVs) are membrane-bound nanoparticles that can be generated by an endosomal sorting complex required for transport-dependent multivesicular body formation or by outward budding of the plasma membrane [15]. EV cargo changes under conditions of health vs. disease [16–18] and function in cell-cell communication [19]. EVs have multiple immunoregulatory functions, and it has been demonstrated in type 1 diabetes that cytokine-induced stress results in the release of EV cargo with inflammatory properties that promote B-cell autoimmunity [16]. Moreover, changes in small EV cargo in the context of beta cell proinflammatory cytokine exposure activate the CXCL10/CXCR3 pathway that increases beta cell dysfunction [20] while lymphocyte-derived exosomes containing the microRNAs miR-142-3p, miR-142-5p and miR-155 promote beta cell death [21].

In cancer cells, PD-L1 expression occurs on EVs to allow tumour cells to evade the immune system [22]. However, the presence of PD-L1 in beta cell EVs has not been described. We hypothesized that islet beta cell inflammation leads to upregulation of PD-L1 in islet beta cell EVs and that, like cellular PD-L1, this beta cell EV PD-L1 may promote beta cell survival in type 1 diabetes.

## Methods

### Cell Culture

Rat insulinoma cells, INS-1 823/13 (INS-1, RRID:CVCL_7226) [23] originally obtained from Chris Newgard (Duke University), NIT-1 insulinoma cell line (RRID:CVCL_3561) provided by Erica Cai (Indiana Biosciences Research Institute) [24] and EndoC-βH1 cells (RRID:CVCL_L909) [25] purchased from Human Cell Design, Toulouse, France were cultured as previously described but containing 10% EV-depleted FBS (see details in the electronic supplementary material [ESM] Method) [18, 26]. Human islets, obtained either through the Integrated Islet Distribution Program or Alberta Islet core (n=8, ESM Human Islet checklist), were cultured as described [18]. To model the inflammatory milieu of type 1 diabetes, cells or human islets were exposed to either 2000 U/ml IFN-α or 100 ng/mL human IFN-□ or 5 ng/mL human IL1-β or cytokine mix of IFN-gamma and IL1-β for 24 h. The list of cytokines is provided as ESM Table 1. Approximately 1×10^7^ INS-1 cells, 8×10^6^ EndoC-βH1 cells and 500 human islet equivalents (IEQs) from the same donor or cell passage were seeded and used for each condition for all experiments unless otherwise stated.

**Table 1.**
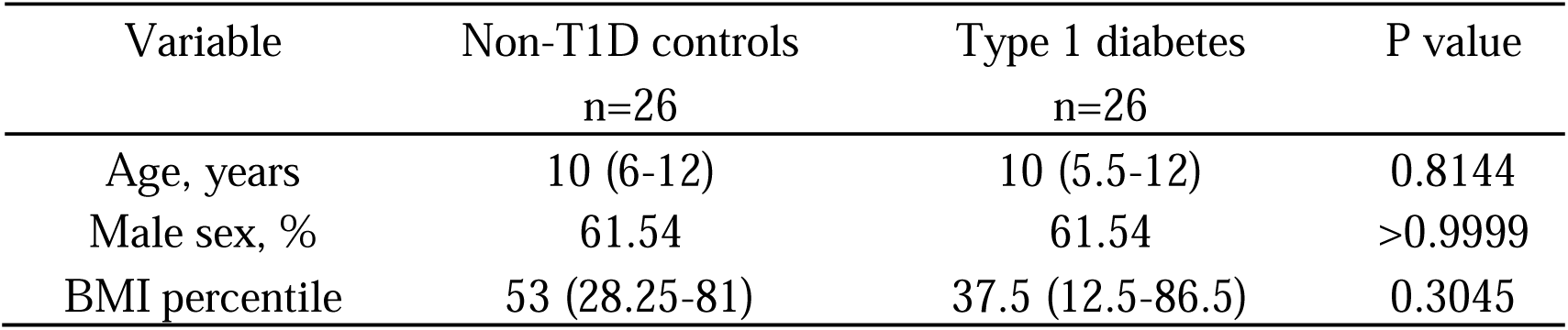
Demographic characteristics of study participants with new-onset clinical type 1 diabetes.

| Variable | Non-T1D controls<br>n=26 | Type 1 diabetes<br>n=26 | P value |
| --- | --- | --- | --- |
| Age, years | 10 (6-12) | 10 (5.5-12) | 0.8144 |
| Male sex, % | 61.54 | 61.54 | >0.9999 |
| BMI percentile | 53 (28.25-81) | 37.5 (12.5-86.5) | 0.3045 |

### EV isolation

Small EVs from cells were isolated from cell culture supernatant using ultracentrifugation, and from human islet medium or plasma using size exclusion chromatography (SEC) as described in ESM Methods [18]. Presence of EVs was verified using nanoparticle tracking analysis (NTA), transmission electron microscopy (TEM) and immunoblot as described in the ESM Methods.

### Immunofluorescence

INS-1 or EndoC-βH1 cells were seeded and allowed to attach overnight in a Millicell EZ slide (Millipore Sigma, Cat#C86024). Adherent cells were fixed with 4% paraformaldehyde for 10 minutes and blocked with 2% BSA and 0.5% Triton X-100 in PBS for 45 minutes. Cells were incubated with primary antibodies against PD-L1 and CD63/CD9/CD81 (ESM Table 2) overnight at 4 C, followed by incubation with secondary antibodies (ESM Table 2). Nuclear staining was performed with Vectasheild mounting medium with DAPI (Vector Laboratories Cat#H-1200-10) and a confocal microscope (LSM800, Carl Zeiss, IL, USA) was used for image analysis.

**Table 2.** Demographic characteristics of nondiabetic study participants with islet autoantibody positivity.

| Variable | Aab- controls<br>n=20 | Aab+<br>n=16 | P value |
| --- | --- | --- | --- |
| Age, years | 12.5 (8-17) | 11.5 (8-28) | 0.9386 |
| Male sex, % | 70 | 81.25 | 0.7003 |
| BMI percentile | 20.15 (18.2-25.4) | 17.5 (16.3-20.2) | 0.1577 |

*Results are displayed as median (interquartile range). BMI- body mass index.*

### Surface EV PD-L1 analysis

PD-L1 surface staining was performed using ExoView® Exosome Human Tetraspanin Kit (Unchained Labs Cat#251-1044, EV-TETRA-C,) or via flow cytometry (see ESM Methods). ExoView® was performed on chips printed with capture antibodies for CD63, CD81 and CD9 according to the manufacturer’s protocol. Briefly, 1 x 10^8^ -5 x 10^8^ EVs/ml was diluted using sample incubation buffer and 50-75 µl of sample were added to the pre-scanned chips and incubated overnight at RT. After incubation, chips were washed and incubated with anti CD9 (CF488, 1:500), anti CD81 (CF555, 1:500) and anti-PD-L1 (FAB1562R, 1:100) for 1 hr. Following washing, the chips were imaged using ExoView R200 automated imager and analysed using ExoView data analysis software.

### PD-1/PD-L1 binding assay

Homogeneous time-resolved fluorescence (HTRF) PD-1/PD-L1 Binding Assay Kit (Cisbio Bioassays SAS, France, Cat#64PD1PEG) was used to test the binding of EV PD-L1 from EndoC-βH1, wild-type HEK293 cells and HEK293 cells overexpressing PD-L1. The detailed protocol is described in the ESM Methods.

### Human samples

Deidentified randomly collected plasma from 26 paediatric subjects with recent-onset type 1 diabetes and from 26 age-, gender-, and BMI-matched non-diabetic controls were obtained from the IU biorepository at Indiana University School of Medicine. For the analysis of soluble PD-L1 34 samples (17 samples from each cohort) were assayed. Deidentified randomly collected plasma from 16 individuals positive for islet autoantibodies and from 20 autoantibody negative nondiabetic controls were also obtained from the IU biorepository at Indiana University School of Medicine. Informed consent and assent as possible were obtained from all patients. Collections were approved by the Indiana University School of Institutional Review Board and reported investigations have been carried out in accordance with the principles of the Declaration of Helsinki as revised in 2008.

### Treatment of CD8 T cells with the EVs

Splenocytes of NOD/ShiLtJ mice (The Jackson laboratories, Cat#001976) were isolated aseptically as described in ESM Methods. To assess T cell-mediated immune activation, 96-well round bottom plates (Greiner Bio One, Cat#650180) were coated with anti-CD3 (0.5 μg/ml, BioLegend Cat# 100202, RRID:AB_312659) and anti-CD28 (1 μg/ml, BD Biosciences Cat# 553294, RRID:AB_394763) antibodies overnight.

Splenic cells (1-1.5 x 10^6^) were cultured with purified NIT-1 wild-type EVs or PD-L1 overexpressing EVs (see ESM Methods) with or without anti-mouse PD-L1 blocking antibody (Bio X Cell Cat# BE0101, RRID:AB_10949073) or Rat IgG isotype control (Bio X Cell Cat# BE0090, RRID:AB_1107780). Cell Trace Violet (CTV)-labelled cells were analysed by flow cytometry at 24 hr, 48 hr, and 72 hr timepoints to assess CD8 T cell proliferation. Cells were also stained with eFluor 780 fixable viability dye (eBiosciences, Cat#65-0865-18) and analysed for activation-linked T-cell surface markers using primary antibodies listed in ESM Table 2. Culture supernatant was collected and stored at −80 °C for cytokine analysis. Samples were recorded on Attune NxT Flow Cytometer (Thermo Fisher Scientific, Waltham, MA, USA), and data were analysed using FlowJo™ v10 Software (BD Biosciences).

### ELISA

Human islet EV, plasma EV and soluble PD-L1 levels were measured using U-plex Human PD-L1 kit (epitope 1) (Mesoscale Discovery). Plasma C-peptide was measured the TOSOH immunoassay (TOSOH Biosciences). EV-splenocyte coculture supernatants were assayed for Mouse Granzyme-B DuoSet ELISA (R&D Systems, Cat#DY1865-05) and IFNγ ELISAs antibody (BD Biosciences). ELISAs were performed according to manufacturer’s protocol and read on a SpectraMax M2 microplate reader (Molecular Devices). All samples were tested in duplicate.

### Statistics

Data were analysed with GraphPad Prism 10.0.3 software for MacOS, GraphPad Software, Boston, MA, USA. Significance was assessed by a two-tailed Student’s t test or Mann–Whitney U test (for nonparametric distributions). Pearsons’ correlation analyses were used to measure monotonic relationships. *P*-values were considered statistically significant when p value<0.05. Data are presented as mean ± SEM.

## Results

### PD-L1 is present in beta cell EVs

To test the idea that PD-L1 may be intracellularly sorted into EV cargo, immunofluorescence staining of INS-1 and EndoC-βH1 cells was performed to characterize PD-L1 co-localization with tetraspanin-associated surface markers, CD63, CD81, or CD9. Here, we observed colocalization of PD-L1, CD63, CD9 or CD81 within cells, supporting the concept that PD-L1 may be sorted with EV cargo **(Fig. 1a, ESM Fig 1a)**. To verify the presence of PD-L1 in beta cell EVs, small EVs were isolated from INS1 rat insulinoma cells, EndoC-βH1 human beta cells, and human islets. EV isolation and purity were validated using TEM **(Fig. 1b),** immunoblot for characteristic EV proteins and calreticulin to ensure depletion of cell debris **(Fig. 1c),** and NTA **(Fig. 2j-l).** The mean diameter of the vesicles measured by TEM and NTA showed values in the range of 50 to 200 nm (majority peaking at 94 nm), consistent with small EVs. Immunoblot analysis confirmed the presence of PD-L1 in EVs, in association with CD9 and CD63, and absence of calreticulin, under control conditions **(Fig. 1c)**.

**Fig. 1.**
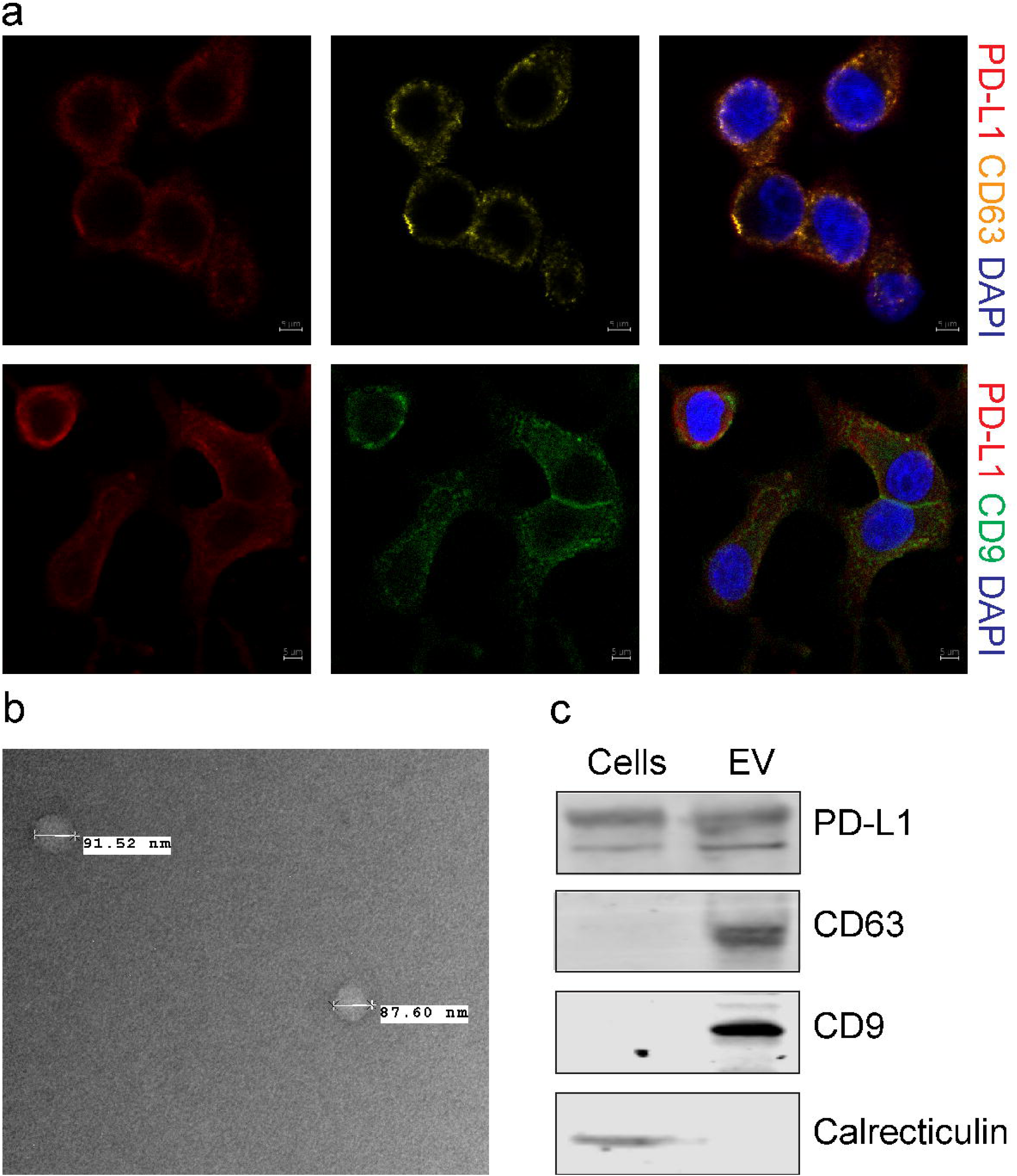
PD-L1 is present in beta cell EVs. **(a)** Confocal microscopy of PD-L1 (red) and tetraspanin-associated surface markers CD63 (yellow), and CD9 (green) in INS-1 cells. **(b**) TEM analysis of INS-1 cell-derived EVs. TEM images display representative data from INS-1 EVs. Scale bars, 500 nm. **(c)** Immunoblot of INS1 cells and EVs for PD-L1, EV markers (CD63 and CD9) and calreticulin (reflects cellular contamination). n=3.

**Fig. 2.**
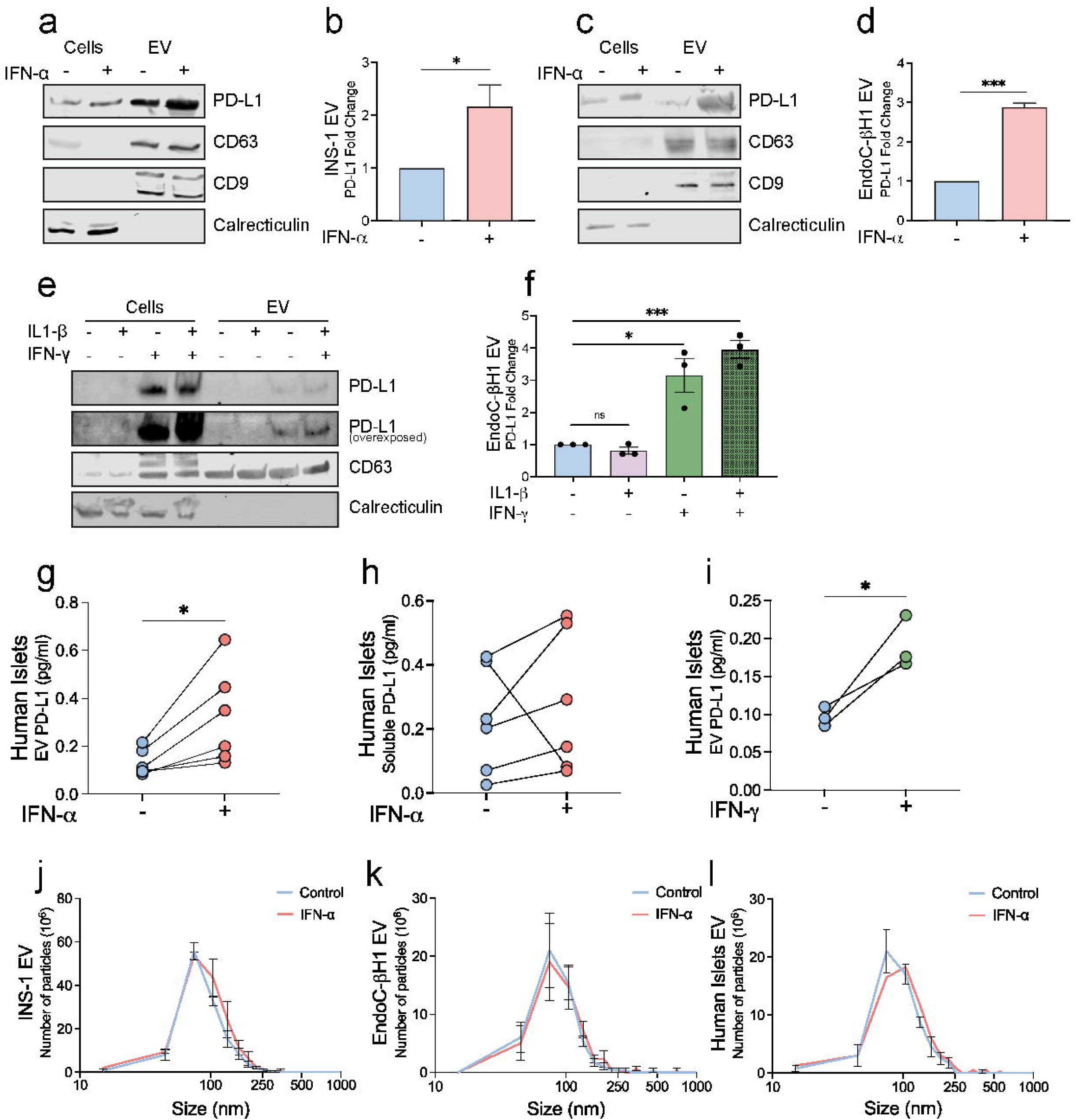
IFN-α or IFN-0 increase beta cell EV PD-L1 cargo. **(a-d)** Immunoblot of **(a-b)** INS1 and **(c-d)** EndoC-βH1 cells and EVs +/-24-h IFN-α for PD-L1, EV markers (CD63 and CD9) and calreticulin. **(b, d)** Quantification of EV PD-L1 protein expression was performed using densitometry and normalized to total protein content (Ponceau S stain) from **(b)** INS-1 and **(d)** EndoC-βH1 EVs. **(e)** Immunoblot of EndoC-βH1 cells and EVs +/-24-h IL1-β, IFN-□ or IL1-β and IFN-□ for PD-L1, CD63 and calreticulin. **(f)** Quantification of EV PD-L1 protein expression was performed using densitometry and normalized to total protein content (Ponceau S stain). **(g)** PD-L1 in human islet EVs +/-24-h IFN-α as measured using ELISA. **(h)** Human islet soluble PD-L1 +/-24-h IFN-α as measured using ELISA. **(i)** PD-L1 in human islet EVs +/-24-h IFN-□ as measured using ELISA **(j-l)** NTA showing EV size distribution of **(j)** INS1, **(k)** EndoC-βH1 and **(l)** human islet EVs +/-24-h IFN-α. Blue bars/lines/circles indicated vehicle control, red bar/lines/circles indicate IFN-α treated samples, and green bar/lines/circles indicate IFN-□ treated samples. n=3 for INS-1 and EndoC-βH1 cells; n=3-6 for human islets. Results are displayed as mean ± SEM *, p value<0.05; ***, p value<0.001.

### IFN-α increases beta cell EV PD-L1 cargo

Recent work has shown that IFN-α or IFN-□, but not IL1-β signalling increases beta cell PD-L1 levels in human beta cells [27–29]. Consistent with this, IFN-α led to a 1.5-fold increase in intracellular PD-L1 expression **(Fig 2a, c).** To determine whether this mode of inflammatory stress increases beta cell EV PD-L1, we also isolated small EVs and quantified PD-L1 content using immunoblot. Here, IFN-α exposure induced a more pronounced, two to three-fold increase in small EV PD-L1 **(Fig. 2a-d).** EndoC-βH1 cells also demonstrated robust EV PD-L1 upregulation upon IFN-□ or IFN-g + IL-1b treatment but not with IL1-β alone **(Fig. 2e-f)**.

Testing of human islets showed an approximate two-fold increase in EV PD-L1 levels with IFN-α or IFN-□ exposure **(Fig. 2g, i)**. We observed a similar trend in soluble PD-L1 levels after IFN-α treatment, but overall differences were smaller, and differences did not reach statistical significance **(Fig. 2h)**. Based on these observations, subsequent experiments were performed with IFN-α or IFN-□.

To understand if increases in EV PD-L1 were due to increased release of EVs under IFN-α stimulated conditions, we performed NTA to determine EV quantity and size distribution. No significant difference in either the particle size distribution or concentration were observed after IFN-α treatment **(Fig. 2j-l)**. These findings suggest that IFN-α treatment does not increase EV size or number, but rather leads to an enrichment of PD-L1 cargo in EVs from human islets and β cell lines.

### PD-L1 protein is present on beta cell EV surface

To test if PD-L1 is expressed on the surface of beta cell EVs and thus, has the potential to directly interact with PD-1 on immune cells, INS-1 EVs were isolated using immunoaffinity capture tetraspanin bead pulldown. Flow cytometry staining of intact/unlysed EVs was performed using an Ab that recognizes an epitope in the extracellular domain of PD-L1. Staining verified the presence of PD-L1 on the EV surface with upregulation after IFN-α or IFN-□ treatment **(ESM Fig. 2a-b)**.

Surface presence of PD-L1 protein in different EV subpopulations was tested by single-particle interferometric reflectance imaging to characterize individual EV particles using an ExoView® microarray chip coated with antibody to human tetraspanin-associated EV markers (CD63, CD81 and CD9) from EndoC-βH1 and human islets treated with or without IFN-α or IFN-□. Fluorescence imaging/ interferometry quantification of surface EV PD-L1 positivity on intact EV subpopulations based on presence of membrane tetraspanins was performed. At baseline, CD63, CD81, and CD9 positive EVs showed similar percentages of PD-L1 positivity on the EV surface (∼10-15%) **(Fig. 3a).** 24-h treatment with IFN-α or IFN-□ yielded significant increases in PD-L1 positive EndoC-βH1 EVs within each subpopulation, but this was the most pronounced for CD81+ EVs (∼6-fold increase with IFN-α and ∼10-fold increase with IFN-□) vs. ∼2-4-fold increase for CD9+ and CD63+ EVs respectively **(Fig. 3c-d)**. In human islet EVs, IFN-α or IFN-□ yielded a ∼1.5-fold increase in PD-L1 positive EVs within each subpopulation **(Fig. 3g-h)**.

**Fig. 3.**
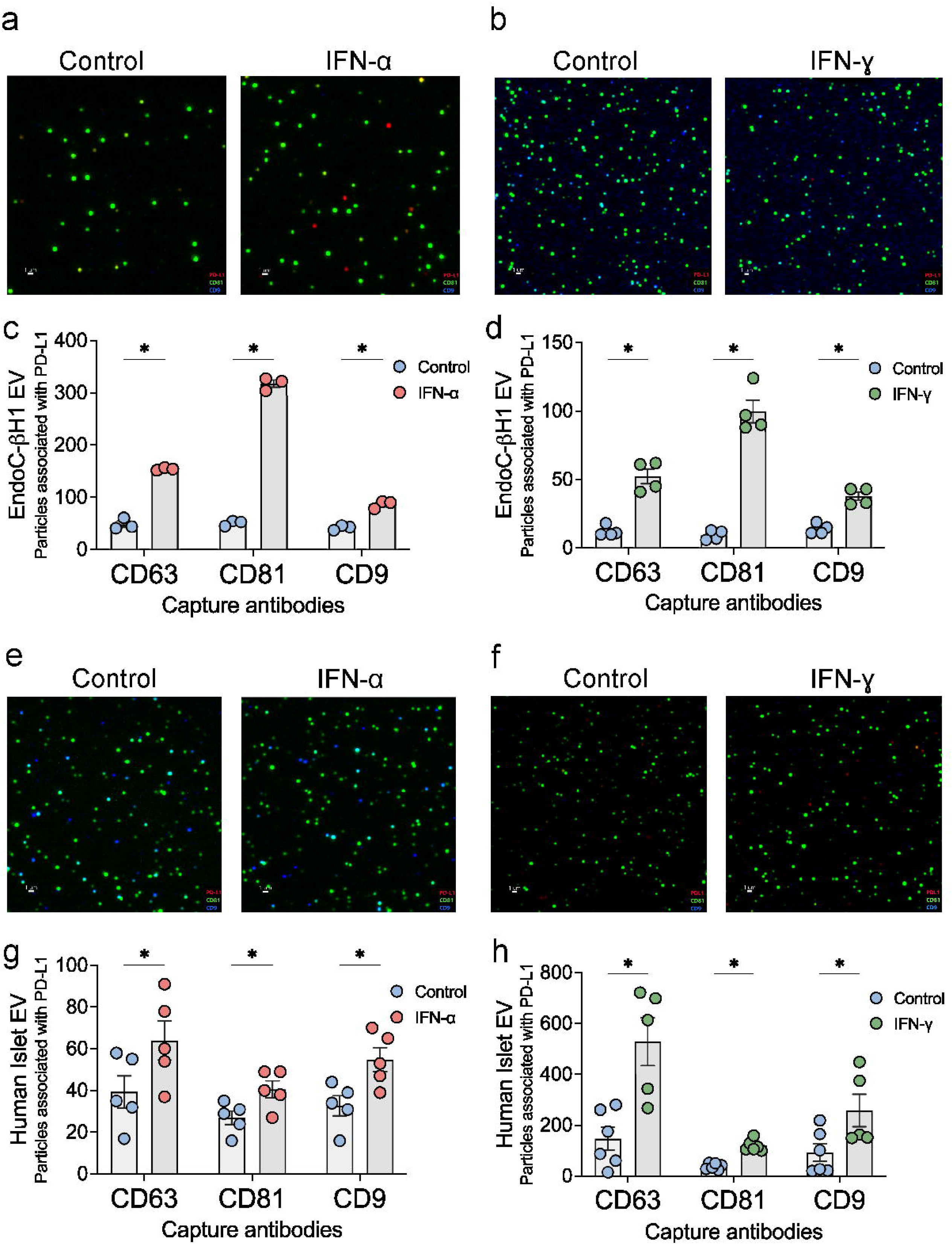
IFN-α or IFN-0 induce PD-L1 expression within tetraspanin-associated EV subpopulations. Capture spot image of **(a-b)** EndoC-βH1 and **(c-d)** total fluorescent particle count of PD-L1 in tetraspanin-associated populations of EndoC-βH1 EVs **(c)** +/-24-h IFN-α or **(d)** +/-24-h IFN-□. Capture spot image of **(e-f)** human islets and **(g-h)** total fluorescent particle count of PD-L1 in tetraspanin-associated populations of human islet EVs **(g)** +/-24-h IFN-α or **(h)** +/-24-h IFN-□. Data represented as mean ± SEM; n=3-4 for EndoC-βH1 EVs and n=4-5 for human islets EVs; *, p value<0.05. Blue circles indicated vehicle control, red circles indicate IFN-α treated samples, and green circles indicate IFN-□ treated samples.

### EV PD-L1 binds PD-1

To test if beta cell EV PD-L1 is capable of binding to PD-1, a PD-1/PD-L1 competitive binding kit was utilized. Decreasing concentrations of EndoC-βH1 EVs were used to test the PD-1 binding and compared to a positive control (PD-L1 standard). A higher concentration of EndoC-βH1 EVs blocked the reagent interaction to interfere with the signal. This interference was reversed with reduced EV concentrations in a dose-dependent manner, confirming binding with PD-L1 on the EV surface (EC50: 0.004±0.002) **(Fig. 4a)**. As a negative control, we performed the binding assay using EVs isolated from HEK293 cells, which do not express PD-L1 **(Fig. 4b).** EVs isolated from HEK293 cells did not effectively block the reagent interaction (EC50: 0.03±0.002) **(Fig. 4c)**. However, EVs isolated from HEK293 cells after transfection with a plasmid to express PD-L1 **(Fig. 4b)** were able to block the interaction in a dose dependent manner with 5.4-fold higher efficacy than the EVs from wild-type HEK293 cells (average EC50: 0.005±0.001) **(Fig. 4c)**. In aggregate, these data confirm that IFN-α increases PD-L1 loading on the beta cell EV surface and that PD-L1 on beta cell EVs can bind PD-1.

**Fig. 4.**
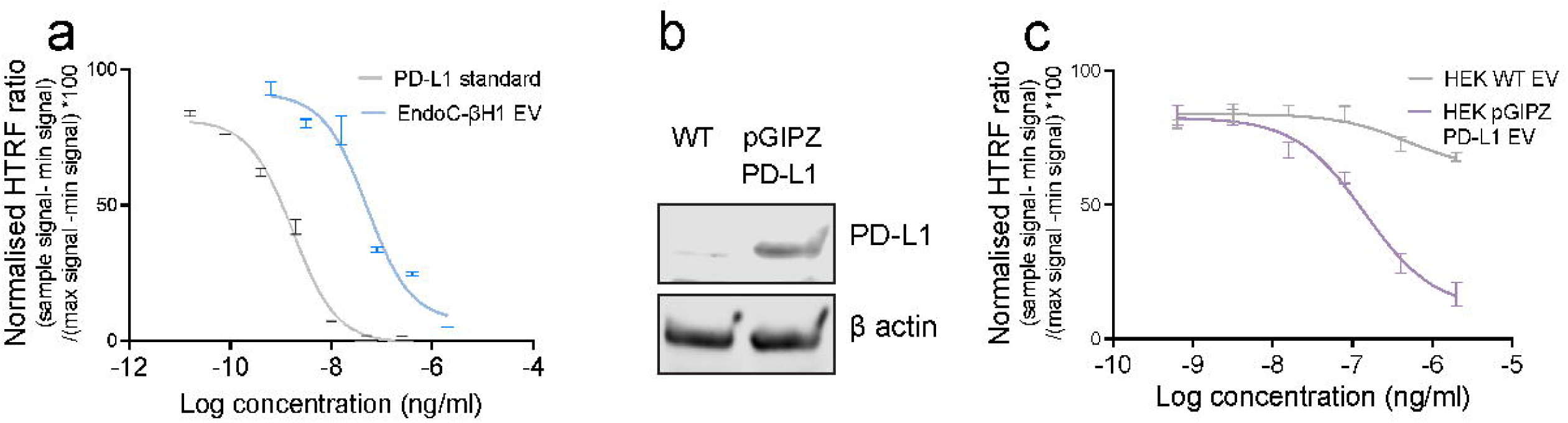
EV PD-L1 can bind PD-1. **(a)** Serial dilution of EndoC-βH1 EV (blue) and PD-L1 standard (grey) as a positive control. **(b)** Immunoblot of HEK293 cells with or without PD-L1 overexpression. β actin is the loading control. **(c)** Serial dilution of EVs from HEK293 wildtype (grey) and HEK293 cells overexpressing PD-L1 (purple). Dose response curves determined by normalized HTRF ratio using non-linear regression analysis. Data represented as mean ± SEM; n=3-4. WT: wildtype.

### EV PD-L1 inhibits proliferation, cytokine production and cytotoxicity of activated murine CD8 T cells in vitro

Next, we investigated if beta cell EV PD-L1 can suppress CD8 T cell function. We exogenously induced PD-L1 expression in NIT-1 cells, which express low levels of endogenous PD-L1, to generate EVs with PD-L1 overexpression (**ESM Fig 3a)**. EVs were isolated from both wildtype (WT) and PD-L1 overexpressing (OE) NIT-1 cells using sequential ultracentrifugation, and EV PD-L1 expression quantified by ELISA **(ESM Fig 3b)**. To assess the impact of EV-associated PD-L1 on immune cells, we isolated splenocytes from NOD mice. Splenocytes were activated with anti-CD3 and anti-CD28 antibodies, then incubated for 72 hours with NIT-1-derived WT or PD-L1^OE^ EVs, and in the presence or absence of a PD-L1 blocking antibody.

Flow cytometric analysis was performed to quantify CD8+ T cell proliferation using CTV dye; activation was quantified with CD69, CD25, or CD44 positivity **(Gating strategy in ESM Fig 3c-d, Results in Fig. 5)**. PD-L1^OE^ EVs significantly inhibited activated CD8+ T cell proliferation **(Fig. 5b, 5f-g)** and activation **(Fig. 5c-e, 5h-m)**. Importantly, pre-treatment of the PD-L1^OE^ EVs with an anti-PD-L1 antibody nearly abolished these inhibitory effects **(Fig. 5b-m)**. PD-L1^OE^ EVs also significantly reduced the cytotoxicity of mouse splenic cells, as demonstrated by decreased levels of IFN-g and granzyme detected in the splenocyte supernatants **(Fig. 5n-o)**. Again, EV pretreatment with anti-PD-L1 antibody abrogated their ability to reduce T cell potential for cytotoxicity **(Fig. 5n-o)**.

**Fig. 5.**
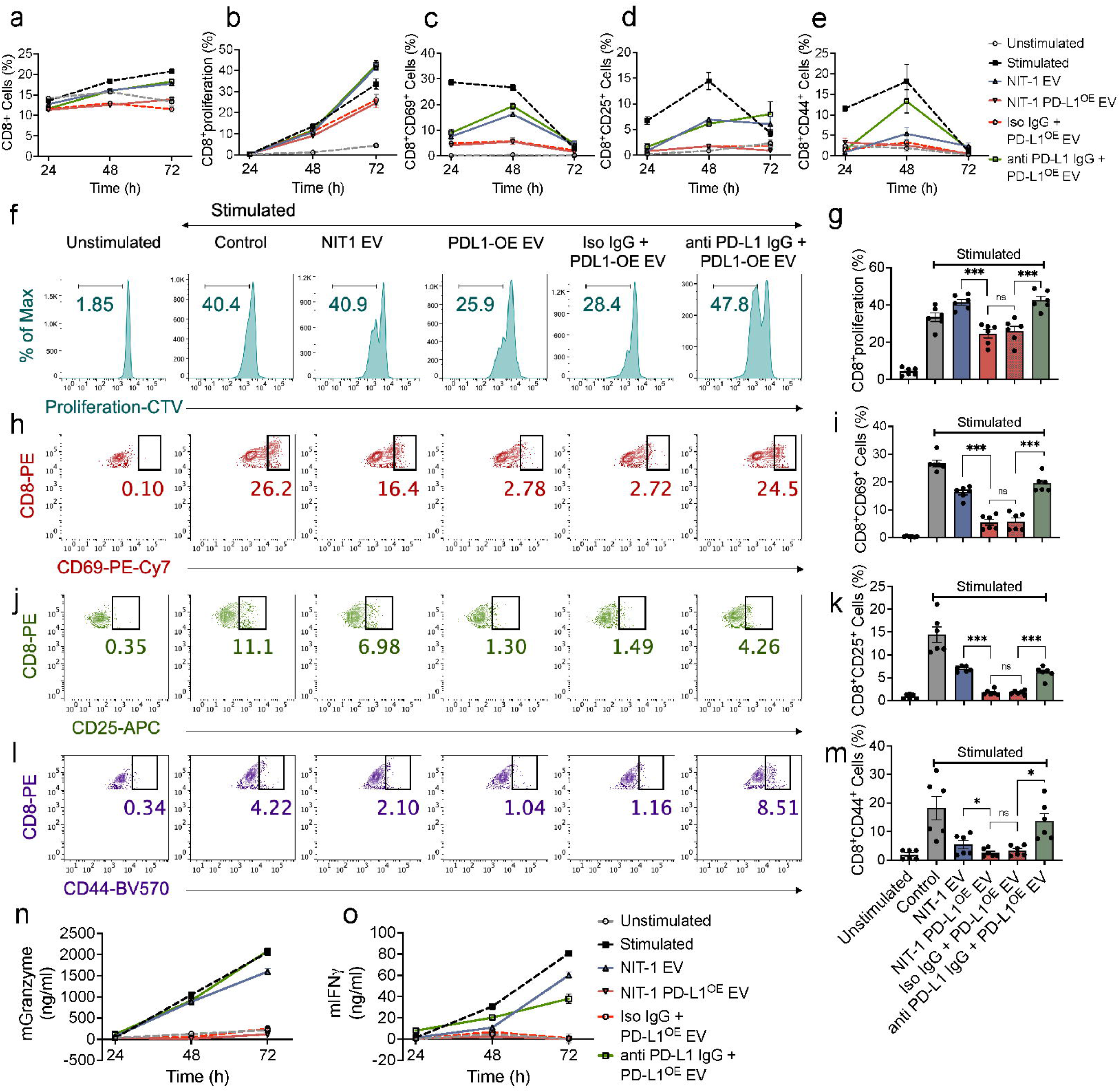
EV PD-L1 inhibits proliferation, cytokine production and cytotoxicity of murine CD8 T cells in vitro. (a-e) Line graph representing the proportion of cells with indicated treatments positive for **(a)** CD8+ T cells **(b)** CD8+ T cells with diluted CTV to assess proliferation and **(c)** CD8+CD69+ expression, **(d)** CD8+CD25+ expression, and **(e)** CD8+CD44+ expression as a measure of CD8+ T cell activation. **(f)** Representative histogram of CTV-labelled NOD CD8 T cells with or without NIT-1 EV treatment at 72 h. **(g)** Bar graphs of proportion of cells with diluted CTV dye. **(h-m)** Representative contour plots and corresponding bar graphs showing proportion of cells from NOD CD8 T cells examined for the surface expression of activation markers: **(h-i)** CD69, **(j-k)** CD25 and **(l-m)** CD44. **(n)** Granzyme secretion and **(o)** IFN-□ secretion from the NOD splenocytes with or without EV treatment as measured by ELISA. Data represented as mean± SEM; n=3; *, p value<0.05; ***, p value<0.001; ns, nonsignificant.

### Plasma EV PD-L1 from children with new-onset type 1 diabetes correlates with residual circulating C-peptide levels

To test the relevance of circulating EV PD-L1 to human type 1 diabetes, we analysed plasma samples from 26 children with recent-onset clinical type 1 diabetes and non-diabetic controls matched by age, sex, and BMI. Demographic characteristics are displayed in Table 1. EVs and non-EV soluble protein were eluted from 500 µl plasma using size exclusion chromatography. We performed NTA on plasma EV samples to assess differences in the overall concentration and size distribution. No difference in total particle number or size distribution existed between the two groups (**Fig. 6a-b**). We next quantified circulating EV PD-L1 and soluble PD-L1 and observed similar soluble and EV PD-L1 levels between children with recent-onset type 1 diabetes and controls **(Fig. 6c-d).** No relationships between plasma EV PD-L1 and age, or sex, or BMI percentile were detected for either group.

**Fig. 6.**
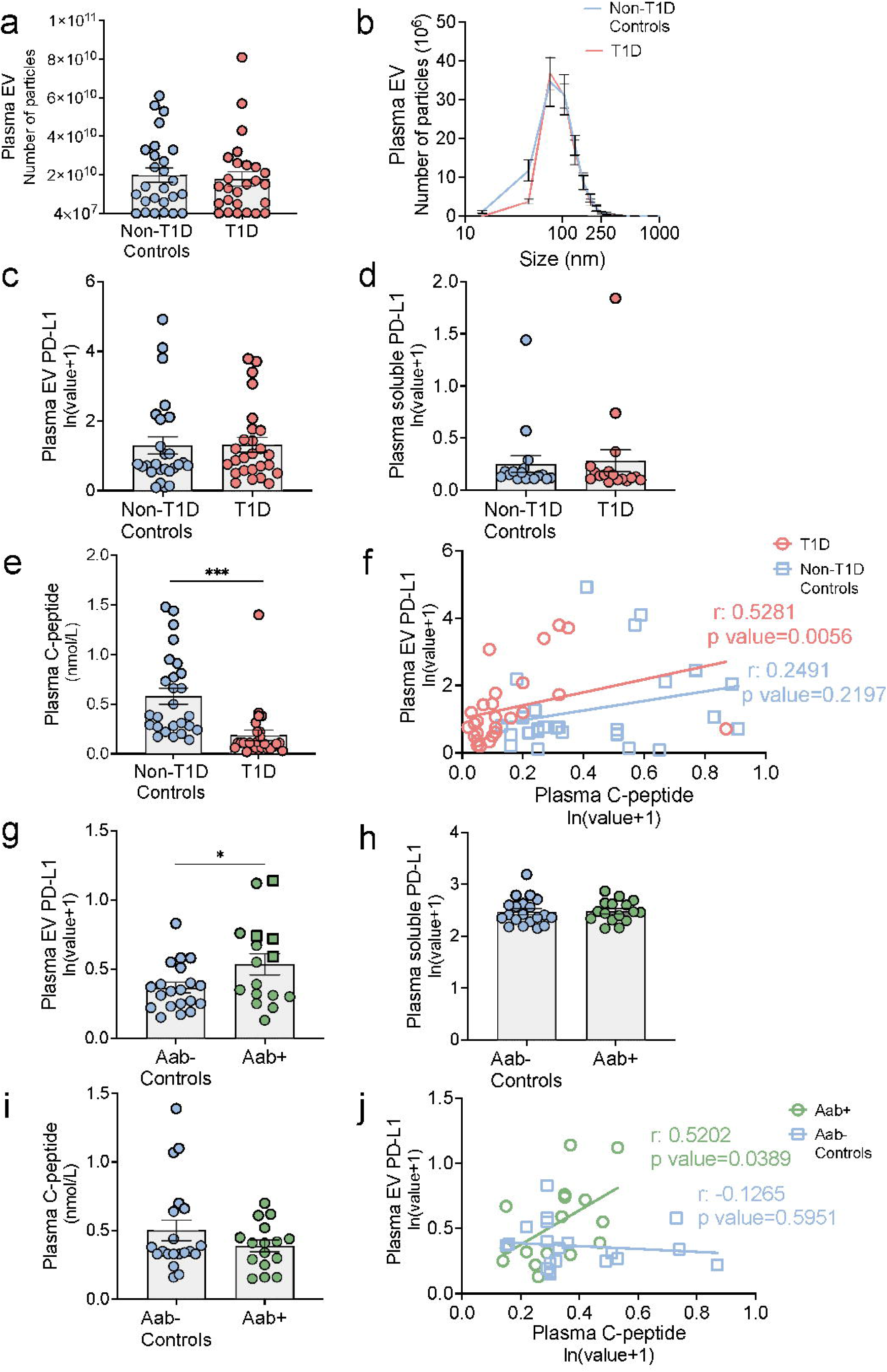
**Plasma EV PD-L1 levels are higher in islet autoantibody positive individuals compared to controls and plasma EV PD-L1 correlates with circulating C-peptide levels in individuals with islet autoantibody positivity or recent-onset clinical type 1 diabetes**. **(a-d)** In plasma from children with recent-onset type 1 diabetes vs. non-diabetic controls: **(a)** Total EV particle number **(b)** EV size distribution **(c)** EV PD-L1 **(d)** Soluble PD-L1 and **(e)** plasma C-peptide levels. **(f)** Linear regression analysis of plasma EV-PD-L1 and plasma C-peptide levels from children with type 1 diabetes (red, T1D) and non-diabetic (blue, non-T1D controls). **(g-k)** In plasma from non-diabetic individuals positive for islet autoantibodies (Aab+) vs islet autoantibody negative (Aab-) controls: **(g)** EV PD-L1 **(h)** Soluble PD-L1 levels **(i)** plasma C-peptide levels, **(j)** Linear regression analysis of plasma EV-PD-L1 and plasma C-peptide levels from islet autoantibody positive (green, Aab+) and autoantibody negative (blue, Aab-controls). Data shown as mean ± SEM, a-c, e-f; n=26/group, d; n=17/group; g-j; n=16/Aab+, n=20/Aab-controls; *, p value<0.05; ***, p value<0.001. Blue circles indicated non-diabetic controls, red circles indicate recent-onset clinical type 1 diabetes samples, green circles indicate non-diabetic islet autoantibody positive samples and green squares indicate single islet autoantibody positive samples.

As expected, plasma C-peptide was significantly reduced in children with type 1 diabetes compared to non-diabetic controls **(Fig. 6e).** However, in children with type 1 diabetes, plasma EV PD-L1 positively correlated with circulating C-peptide (spearman r=0.5281; p value=0.0056) **(Fig. 6f, Red)**, suggesting a link between EV PD-L1 and residual beta cells in type 1 diabetes. This relationship was not present in non-diabetic participants **(Fig. 6f, blue)**.

To evaluate if circulating levels of EV-associated and soluble PD-L1 would vary prior to the development of type 1 diabetes we analysed EV and soluble PD-L1 levels from previously banked plasma samples from 16 nondiabetic individuals with islet autoantibody positivity and 20 autoantibody-negative nondiabetic controls (demographics in Table 2). Here EV PD-L1 levels were elevated in individuals with islet autoantibody positivity compared to controls **(Fig. 6g)**.

Interestingly, this increase was not observed for soluble PD-L1 levels **(Fig. 6h)**. Upon stratification based on the number of islet autoantibodies, we observed that those with a single islet autoantibody exhibited higher levels of plasma EV PD-L1 **(Fig. 6g, green squares)**. We also examined the relationship between circulating C-peptide levels (**Fig. 6i**) and plasma EV PD-L1. Similar to recent-onset clinical type 1 diabetes, a positive correlation was observed between plasma EV PD-L1 and circulating C-peptide (Pearson’s r = 0.5202; p-value =0.0389) in individuals predisposed to type-1 diabetes (**Fig. 6j, green)**. This correlation suggests a link between EV PD-L1 and residual beta cell function in these individuals at-risk for type-1 diabetes. Notably, this relationship was not present in the control group (**Fig. 6j, blue**).

## Discussion

Multiple groups have demonstrated the potential for beta cell PD-L1 to promote beta cell survival in the context of islet inflammation and autoimmunity [30–34]. However, to our knowledge, this is the first description of the presence of PD-L1 in beta cell EVs. Our findings suggest that PD-L1 is present on the surface of small EVs secreted by pancreatic beta cells and EV PD-L1 is upregulated in response to IFN-α or IFN-□ in beta cell lines and primary human islets. EV PD-L1 has the capacity to bind to PD-1 and significantly impairs the proliferation, activation, and cytotoxicity of activated CD8 T cells. Circulating EV PD-L1 is increased in some individuals with islet autoantibody positivity, especially those with single-positive autoantibodies. Positive associations between circulating EV PD-L1 levels and C-peptide in humans with autoantibody positivity or clinical type 1 diabetes support a protective role for circulating EV PD-L1 in this population.

Our previous studies have been shown that IFNs, but not IL-1b or TNF, upregulate PD-L1 expression in human beta cells [27, 29]. IFN-α plays a prominent role in mediating host viral responses and in the autoimmune response during early type 1 diabetes development. A type 1 IFN gene signature is present in individuals during early type 1 diabetes development, with elevated plasma IFN-α. In the human islet microenvironment, IFN-α exposure increases beta cell PD-L1 expression via JAK/STAT-IRF1 signalling. The present studies go a step further, showing that IFN exposure not only increases total cell PD-L1 expression but also increases PD-L1 shuttling into EV cargo in multiple EV subpopulations.

Prior work has suggested that beta cells upregulate PD-L1 to limit destruction by self-reactive immune cells. PD-L1 inhibition in NOD mice increases insulitis progression and diabetes [30, 31] while PD-L1 overexpression prevents immune destruction in a xenograft model [35] and PD-L1 expression is increased in beta cells that survive the immune attack in NOD mice [36]. Engineered artificial EVs overexpressing PD-L1 and Gal-9 proteins were able to reverse new-onset hyperglycemia and delay diabetes onset in this model [37]. Crucially, we show that beta cell PD-L1 on the EV surface can directly bind to PD-1, with functional studies demonstrating the immunomodulatory capacity of beta cell EV PD-L1. These findings suggest that beta cells may employ EV PD-L1 as an additional mechanism to modulate autoreactive immune responses in type 1 diabetes, providing a paracrine signal to amplify immune checkpoint signalling by the beta cell. This possibility is supported by our data in human samples, showing a positive relationship between circulating EV PD-L1 with residual C-peptide in individuals with autoantibody positivity or clinical type 1 diabetes.

Prior work showed variable PD-L1 expression in insulin-containing islets from recent-onset type 1 diabetes donors, correlating with increased CD8 T cell presence [27]. We also observed heterogeneity in circulating EV PD-L1 in autoantibody-positive donors and children with recent-onset type 1 diabetes, with the highest levels in single versus multiple autoantibody-positive individuals. These findings may reflect that heterogeneity in PD-L1 expression is related to timing in the context of the natural history of the disease and in the presence of active autoimmunity. Indeed, elevated plasma EV PD-L1 in nondiabetic islet autoantibody-positive individuals but not in those with recent-onset type 1 diabetes, suggest that changes in circulating PD-L1 may be more prominent earlier in the natural history of the disease, possibly coinciding with the importance of IFN-α during early type 1 diabetes development [38–40]. Alternatively, preferential expression of PD-L1 in insulin-positive cells from donors with type 1 diabetes or autoantibody positivity and correlations of circulating EV PD-L1 with C-peptide could reflect that heterogeneity is linked to a protective effect on beta cell survival or a mechanism to restrain further epitope spreading [27, 41]. Longitudinal studies of autoantibody-positive individuals with larger sample sizes will help to better elucidate these relationships.

EVs are typically categorized into three size-based subpopulations [15]. Small EVs, often termed exosomes, form by fusion of the multivesicular endosome with the plasma membrane, while larger EVs (microvesicles and apoptotic bodies) emanate from plasma membrane blebbing [15]. We specifically examined the association of PD-L1 with small EVs upon IFN exposure, but other studies focusing on PD-L1+ tumour EVs have detected the presence of PD-L1 in microvesicles, albeit at a lower level compared to the levels present in small EVs [22, 42]. An important next question will be the existence of beta cell EV PD-L1 enrichment within EV subtypes and the impact of regulators of beta cell EV biogenesis and cargo sorting on PD-L1 cargo. Interestingly, PD-L1 induction was most strongly linked with CD81+ EVs compared to other tetraspanins in EndoC-βH1 EVs. Recent work has shown that CD81 is upregulated in diabetic conditions, with expression of CD81 increased in stressed human beta cells [43].

Differences in islets vs. beta cells may reflect contributions of other islet cells. Future studies should investigate the role of CD81+ PD-L1+ beta cell EV subpopulations in triggering or repressing signalling cascades important in type 1 diabetes autoimmunity and PD-L1 positivity in EVs emanating from other islet cells.

A limitation for detection of EV PD-L1 from circulation is that islet-derived EVs may be diluted by other circulating EV populations. Emerging strategies to isolate EVs specifically derived from islet beta cells [44, 45], holds potential to enhance the specificity of assays and enable investigators to focus on biomarkers that are directly related to the health and function of these cells. Our observed relationship with residual C-peptide levels in both at-risk individuals and type 1 diabetes is suggestive of a beneficial contribution of islet derived EV PD-L1 in preserving beta cells. The absence of this correlation in non-diabetic individuals may indicate that EV PD-L1 becomes particularly relevant in the context of ongoing autoimmune attack. Future studies should investigate the relationship of longitudinal circulating EV PD-L1 levels and insulin secretion, at different stages of type 1 diabetes disease progression.

EVs can also serve as natural drug delivery vehicles and therapeutic molecules, implying that PD-L1 could be encapsulated within EVs and targeted to specific cell types or tissues, potentially including beta cells or immune cells involved in the autoimmune response [46–48]. A beneficial effect of PD-L1+ EVs on beta cell tolerance could theoretically be harnessed as an intervention to prevent autoimmune beta cell destruction [46]. Further studies will test the significance of beta cell EV PD-L1 transfer to other recipient cells in the progression to type 1 diabetes and establish if beta cell EV PD-L1 transfer indeed represents a feasible target.

In conclusion, our data identify EV PD-L1 as an additional mechanism supporting the effects of the PD-L1/PD-1 axis in type 1 diabetes both at the level of the islet and in circulation. These findings identify a plausible framework for EV PD-L1 binding to PD-1, allowing a mechanistic basis for immune modulation and suggesting that EV PD-L1 could serve as a predictive biomarker during development of type 1 diabetes. Longitudinal studies as individuals progress through disease stages will allow for a better understanding of the heterogeneity in circulating EV PD-L1. Future studies should define the pathophysiological relevance of EV PD-L1 on other surrounding islet and immune cells in the context of type 1 diabetes to further elucidate the potential of EV PD-L1 as an intervention to impact immune cell activity.

## Acknowledgements

The authors thank Anthony Acton and Gabriella Monaco at Indiana University for technical assistance with PD-L1 and C-peptide ELISA, and Nicholas Conoan of the Electron Microscopy Core Facility (EMCF) at the University of Nebraska Medical Center for TEM. The EMCF is supported by state funds from the Nebraska Research Initiative (NRI) and the University of Nebraska Foundation, and institutionally by the Office of the Vice Chancellor for Research. This work also utilized the Indiana University Flow Cytometry Core. The authors thank the members of the Indiana University Melvin and Bren Simon Comprehensive Cancer Center Flow Cytometry Core. The Indiana University Melvin and Bren Simon Comprehensive Cancer Center Flow Cytometry Core is funded in part by NIH, National Cancer Institute (NCI) grant P30 CA082709. This study used core services provided by the Indiana Diabetes Research Center grant P30 DK097512 (to Indiana University School of Medicine).

## Contribution Statement

CR and DTC acquired, analysed and interpreted the data, and drafted the manuscript. SR, AGO, and JX acquired the data and reviewed the manuscript critically for important intellectual content. CEM, JDP, DLE and RGM made substantial contributions to the design of the work and reviewed the manuscript critically for important intellectual content. CR and EKS made substantial contributions to the conception of the work, interpreted data and drafted the manuscript. All the authors approve the version of the manuscript to be published. EKS is the guarantor of this work and, as such, had full access to all the data in the study and takes responsibility for the integrity of the data and accuracy of the data results.

## Data Availability

The datasets generated during and /or analysed during the current study are available from the corresponding author on reasonable request.

## Abbreviations

AAB: Islet-specific autoantibody positive
CTV: Cell-Trace Violet
CXCL10: CXC motif chemokine ligand 10
CXCR3: CXC motif chemokine receptor 3
EV: Extracellular Vesicle
HTRF: Homogeneous time-resolved fluorescence
IEQ: Islet Equivalents
IRF: IFN regulatory factor
JAK: Janus Kinase
NTA: Nanoparticle Tracking Analysis
OE: Overexpressing
PD-L1: Programmed cell death 1 Ligand -1
PD-1: Programmed cell death 1
RT: Room Temperature
SEC: Size Exclusion Chromatography
STAT: Signal transducer and activator of transcription
TEM: Transmission Electron Microscopy
WT: Wild type

