## Supplementary Material for "Beta cell extracellular vesicle PD-L1 as a novel regulator of CD8+ T cell activity and biomarker during the evolution of Type 1 Diabetes"

**Electronic Supplementary Material**

**Manuscript Title**

**Authors and affiliations**

Chaitra Rao[1,2,3,7], Daniel T Cater[1,7], Saptarshi Roy[4], Jerry Xu[1,2,3], Andre De G Olivera[1,2,3], Carmella Evans-Molina[1,2,3,4], Jon D. Piganelli[4], Decio L. Eizirik[5], Raghavendra G. Mirmira[6], Emily K. Sims*[1,2,3]

1 Department of Pediatrics, Indiana University School of Medicine, Indianapolis, IN, USA
2 Center for Diabetes and Metabolic Diseases, Indiana University School of Medicine, Indianapolis, IN, USA
3 Herman B. Wells Center for Pediatric Research, Indiana University School of Medicine, Indianapolis, IN, USA
4 Department of Medicine, Indiana University School of Medicine, Indianapolis, IN, USA
5 ULB Center for Diabetes Research, Medical Faculty, Université Libre de Bruxelles (ULB), Brussels, Belgium.
6 Department of Medicine and the Kovler Diabetes Center, The University of Chicago, Chicago, IL, USA
7 Authors contributed equally to this work.
* Corresponding author

**Corresponding author**

Emily K. Sims M.D. M.S
Associate Professor of Pediatrics,
Indiana University School of Medicine
Center for Diabetes and Metabolic Diseases

635 Barnhill Drive, MS 2031
Indianapolis, IN 46202
317-274-4130 tel


**Content**

ESM Methods

ESM Tables

ESM Figures

References

Human islet checklist

**Abbreviations:**

| | |
|---|---|
| APC | Allophycocyanin |
| EV | Extracellular Vesicle |
| ECM | Extracellular Matrix |
| HTRF | Homogeneous time-resolved fluorescence |
| IEQ | Islet Equivalents |
| NTA | Nanoparticle Tracking Analysis |
| PD-L1 | Programmed death Ligand -1 |
| PD-1 | Programmed death -1 |
| SEC | Size Exclusion Chromatography |
| TEM | Transmission Electron Microscopy |

**ESM Methods:**

**Cell Culture**: INS-1 832/13 cells were cultured in RPMI supplemented with 10% EV-depleted FBS (prepared by ultracentrifugation at 120,000 *g* for 20 h at 4°C followed by filtration [0.22 µm PES filter]), 10 mm HEPES (Gibco, Thermo Fisher Scientific), 100 units/ml penicillin + 100 µg/ml streptomycin (Corning) as described [1, 2]. EndoC-βH1 cells were cultured in low glucose DMEM containing 5.5 mmol/l glucose 2% BSA fraction V, fatty acid free (Roche, Basel, Switzerland), 50 µmol/l 2-mercaptoethanol (Sigma-Aldrich), 10 mmol/l nicotinamide (Calbiochem), 5.5 µg/ml transferrin (Sigma-Aldrich), 6.7 ng/ml sodium selenite (Sigma-Aldrich), 100 U/ml penicillin + 100 µg/ml streptomycin (Corning), in ECM/fibronectin-coated plates. NIT-1 cells were maintained in DMEM (Gibco, Cat#10313039), supplemented with 10% EV-depeleted FBS (Gibco), GlutaMAX (Gibco), and penicillin/streptomycin (Corning). Cells were grown in a 37°C incubator with 5% CO2 and were free of mycoplasma, as determined monthly using the MycoAlert Mycoplasma Detection kit (Lonza). Available cells or islet preparations were not randomised but divided evenly for comparison of control and cytokine treatments among islets from the same donor or cells from the same parent cells. All the experimental replicates correspond to different donors, and the donor characteristics are detailed in ESM Human Islet checklist.

**EV isolation:** For cell lines, 10-15 ml of supernatant was centrifuged at 800 *g* for 10 minutes. The supernatant fraction was centrifuged at 2,000 *g,* passed through a 0.22 µm filter (Merck Millipore, Cat#GSWP02500) and was centrifuged at 10,000 *g* for 1.5-h at 4°C to pellet small EVs. Total EV protein concentrations were determined by using a BCA protein assay kit (Thermo Fisher Scientific, Cat# 23225). EVs were isolated from human islet medium and 500 µl plasma using size exclusion chromatography (SEC). Samples were centrifuged at 2,000 *g* for 10 minutes and ultrafiltered using 1 µm filter (Cytivia, GE Healthcare distributor, Cat#6780-2510). After rinsing the qEV columns (IZON Science, Cat#ICO-70) with 0.22 µm filtered 1X PBS, 500 µl of the sample was applied on top of a column and 0.5 ml fractions were collected in 1.5 ml tubes. Four EV-rich fractions (6-9) and six soluble protein rich fractions (13-18) were pooled and analysed for EV purity.

**Nanoparticle Tracking Analysis:** EV enriched samples were analysed for concentration and size distribution with dynamic light scattering using a ZetaView instrument (ParticleMetrix, GmbH, Ammersee, Germany) with 100 nm microspheres as particle size standards as described previously [1]. Samples were diluted in 0.22 µm filtered PBS to a concentration within the manufacturer's recommendations. 1 ml diluted sample was loaded into the flow cell and was recorded for 55 seconds. Particle sizes and numbers was analysed at 11 positions per sample and calculated as the mean of the results with ZetaView Analyse software (ParticleMetrix).

**Transmission Electron Microscopy (TEM):** Samples were spotted onto formvar/silicon monoxide coated 200 mesh copper grids (Ted Pella Inc. Redding, CA). Grids were glow discharged for 60 seconds at 20µA with a GloQube glow discharge unit (Quorum Technologies, East Sussex, UK) prior to use. Samples were negatively stained with NanoVan (Nanoprobes, New York, NY) and examined on a Tecnai G2 Spirit TWIN (FEI, Hillsboro, OR) operating at an accelerating voltage of 80kV. Images were acquired digitally with an AMT (Woburn, MA) digital imaging system.

**Immunoblot**: Proteins were extracted from cell lysate or from pelleted EV fractions using 1X RIPA buffer (Thermo Fisher Scientific Cat#89900) with protease and phosphatase inhibitor cocktail (Thermo Fisher Scientific Cat#78441). Samples were diluted using 1X sample buffer (LI-COR Biosciences Cat#928-40004) with 2-Mercaptoethanol (Thermo Fisher Scientific, Cat# 21985023). The proteins were separated using 4-20% SDS-PAGE precast gel (Bio-Rad, Cat#4561094, Cat##4561096) and transferred to activated PVDF membrane. Following blocking with Odyssey blocking buffer (LICOR-Biosciences Cat#927-50003) for 45 minutes, the membrane was incubated with primary antibodies (ESM Table 2) overnight at 4 °C. The membrane was then incubated with IRDye 800CW and 680RD secondary antibodies (ESM Table 2) and imaged on the Odyssey CLX Scanner (LI-COR Biosciences, Lincoln, NE, USA).

**Flow cytometry:** Briefly $1x10^7$ of streptavidin beads (Thermo Fisher Scientific Cat#10608D,) were diluted in 1 ml PBS/1% BSA and incubated with biotinylated CD9 or CD63 antibody at room temperature for 1 hr according to the manufacturer's protocol. Pelleted INS-1 EVs +/- IFN treatment were co-incubated with antibody-coupled magnetic beads overnight at 4°C on a sample mixer. Samples were stained with APC anti-PD-L1 and Exo-FITC (System Biosciences Cat#EXOFLOW800A-1) and washed with 1 ml PBS/1% BSA to remove the unbound antibodies. Samples were run on BD LSR Fortessa (X-20, BD Biosciences, CA, USA) for 2 min and data were analysed using FlowJo™ v10.8 Software (BD Biosciences). The list of antibodies is provided as ESM Table 2.

**Generation of PD-L1 overexpressing cell lines:** HEK293 cells were transiently transfected with pGIPZ-PD-L1-EGFP DNA (RRID: Addgene_120933) using Lipofectamine 3000 (Thermo Fisher Scientific Cat#L3000001) for 72 hours. To generate PD-L1 overexpression in NIT-1 cells, murine PD-L1 plasmid pGIPZ-mPDL1 (RRID: Addgene_121488) was packaged into lentiviral particles using HEK 293T cells co-transfected with viral packaging plasmids [3]. Lentiviral supernatants were harvested after 72 h, and NIT-1 cells were infected with filtered lentivirus and selected using puromycin (5 µg/ml, InvivoGen, Cat#ant-pr-1,). Presence of PD-L1 overexpression in the cells was confirmed either by immunoblot. Presence of EV PD-L1 from NIT-1 cells with or without PD-L1 overexpression was detected using the Mouse PD-L1 DuoSet ELISA kit (R&D Systems, Cat#DY1019-05).

**PD-1/PD-L1 binding assay:** The PD1/PD-L1 binding assays were performed in white 96-well low volume plates (Revvity Health Sciences Inc, Cat#66PL96025) with a final volume of 20 µl comprising 2 µl of diluted EndoC-βH1 or HEK293 EVs/standard, 4 µl of Tag1-PD-L1 (5 nM) and 4 µl of Tag2-PD-1 (50 nM). Following 10 min of incubation, 10 µL of pre-mixed anti-Tag1-Europium and anti-Tag2-XL665 detection reagents were added. HTRF signal was measured after 2 hours using a microplate reader (SpectraMax iD5, Molecular Devices, CA, USA) and the measurement conditions were set up in the SoftMax® Pro software. Results were analysed with a two-wavelength signal ratio: (intensity (665 nm)/intensity (620 nm)) $\times 10^4$ (HTRF Ratio). The normalized HTRF ratio was calculated in accordance with the guidelines provided by the manufacturer which is as follows: ((sample signal) − (min signal))/ ((max signal) − (min signal)) ×100, where 'max signal' is the signal ratio with tagged PD-1/PD-L1 proteins and 'min signal' the signal ratio without tagged PD-1 protein.

**Splenocyte isolation:** Splenocytes of seven–to-twelve-week-old NOD/ShiLtJ mice (The Jackson laboratories, Cat#001976), were isolated aseptically. Spleens were homogenized in MACS buffer (1xPBS, 0.5% BSA and 2m EDTA), filtered through a 40 µm strainer (Fisherbrand, Cat#22363547) and treated with RBC lysis buffer (Sigma, Cat#R7757-100). The cell suspension was washed with MACS buffer and centrifuged at 500 g for 5 minutes. Cells were then stained with CellTrace Violet (CTV) (Invitrogen, Cat#C10094) as follows: a 5 mM stock solution was prepared and diluted 1:2000 in 1X PBS to create a 2.5 µM working solution. After warming the diluted CTV for 5 minutes at 37°C, cells were resuspended at $1 \times 10^6$ cells/mL and incubated at 37°C in a darkened water bath for 10 minutes. The reaction was stopped by adding 5 volumes of cold medium with 10% FBS, followed by resuspension in 5 mL splenocyte medium, containing RPMI medium (Gibco), 10% heat-inactivated FBS (Gibco, Thermo Fisher Scientific, Cat#S11550H, Lot-A20005) and supplemented to a final concentration with L-glutamine (Thermo Fisher Scientific, 2 mM), penicillin (Corning, 50 U/ml), streptomycin (Corning, 50 µg/ml), 2-mercaptoethanol (Thermo Fisher Scientific, 50 µM), 1% HEPES (1M Thermo Fisher Scientific, Cat#15630080), 1% Sodium pyruvate (Thermo Fisher Scientific, 100mM), 1%NEAA (Thermo Fisher Scientific, Cat#11140-050) and 0.7% 2-Mercaptoethanol (Sigma, Cat#M6250).

ESM Tables

| Reagent | Source | Company | Cat. # | Concentration |
|---|---|---|---|---|
| IFN-Alpha 2a | Human | pbl assay sci, Pestka Biomedical Laboratories | 11100 | 2000 U/ml |
| IFN-gamma | Human | R&D systems, Minneapolis, MN, USA | 285IF100 | 100 ng/ml |
| IL1-β | Human | R&D systems, Minneapolis, MN, USA | 201LB005 | 5 ng/ml |
| IFN-Alpha 1 | Rat | pbl assay sci, Pestka Biomedical Laboratories | 13101-1 | 2000 U/ml |
| IFN-gamma | Rat | R&D systems, Minneapolis, MN, USA | 585IF100 | 100 ng/ml |

**ESM Table 1: List of reagents used in the present study**

| Antibody | Source | Company | Cat. # | RRID | Dilution | Application |
|---|---|---|---|---|---|---|
| PD-L1 | Mouse, Rat | Proteintech | 17952-1-AP | AB_10597552 | 1:1000 | WB, IF |
| PD-L1 | Human | Cell Signaling | 13684 | AB_2687655 | 1:1000 | WB |
| PD-L1 | Human | R&D Systems | FAB1562R | | 1:200 | IF, Flow |
| PD-L1 | Rat | BioLegend | 124311 | AB_10612935 | 1:200 | FCM |
| CD63 | Rat, Human | Antibodies Online | ABIN144001 | | 1:1000, 1:200 | WB, IF |
| CD9 | Rat, Human | Proteintech | 20597-1-AP | AB_2878706 | 1:1000, 1:200 | WB, IF |
| PE-CD81 | Human | BD Pharmingen | 555676 | AB_396029 | 1:200 | IF |
| Calrecticulin | Rat, Human | Proteintech | 27298-1-AP | AB_2880835 | 1:1000 | WB |
| β-actin | Mouse, Human | Cell Signaling | 3700 | AB_2242334 | 1:2000 | WB |
| Biotin-CD9 | Rat, Human | Novus | SN4C3-3A2 | AB_10007929 | | FCM |
| Biotin-CD63 | Rat | Biorbyt ltd | orb452831 | | | FCM |
| Biotin-CD63 | Human | Novus | 42225B | AB_2884028 | | FCM |
| FITC CD45 | Rat | Thermo Fisher Scientific | 11-0451-82 | AB_465050 | | FCM |
| PE CD8a | Rat | BioLegend | 162303 | AB_2894434 | | FCM |
| APC CD25 | Rat | BioLegend | 101910 | AB_2280288 | | FCM |
| BV605 CD44 | Rat | BioLegend | 103047 | AB_2562451 | | FCM |
| PE/Cyanine7 CD69 | Armenian Hamster | BioLegend | 104512 | AB_493564 | | FCM |
| IRDye 800CW anti-Rabbit IgG | Goat | Licor Biosciences | 926-32211 | AB_621843 | 1:10000 | WB |
| IRDye 680RD anti-Mouse IgG | Mouse | Licor Biosciences | 926-68072 | AB_10953628 | 1:10000 | WB |
| IRDye 680RD anti-Goat IgG | Donkey | Licor Biosciences | 926-68074 | AB_10956736 | 1:10000 | WB |
| anti-Rabbit IgG (H+L) Texas Red | Goat | Thermo Fisher Scientific | T-2767 | AB_2556776 | 1:1000 | IF |
| anti-Goat IgG (H+L) Alexa Fluor™ 647 | Donkey | Thermo Fisher Scientific | A-21447 | AB_2535864 | 1:1000 | IF |
| anti-Mouse IgG (H+L), Alexa Fluor™ 488 | Donkey | Thermo Fisher Scientific | R37114 | AB_2556542 | 1:1000 | IF |

**ESM Table 2: List of antibodies used in the present study and conditions of use.**

ESM Figures

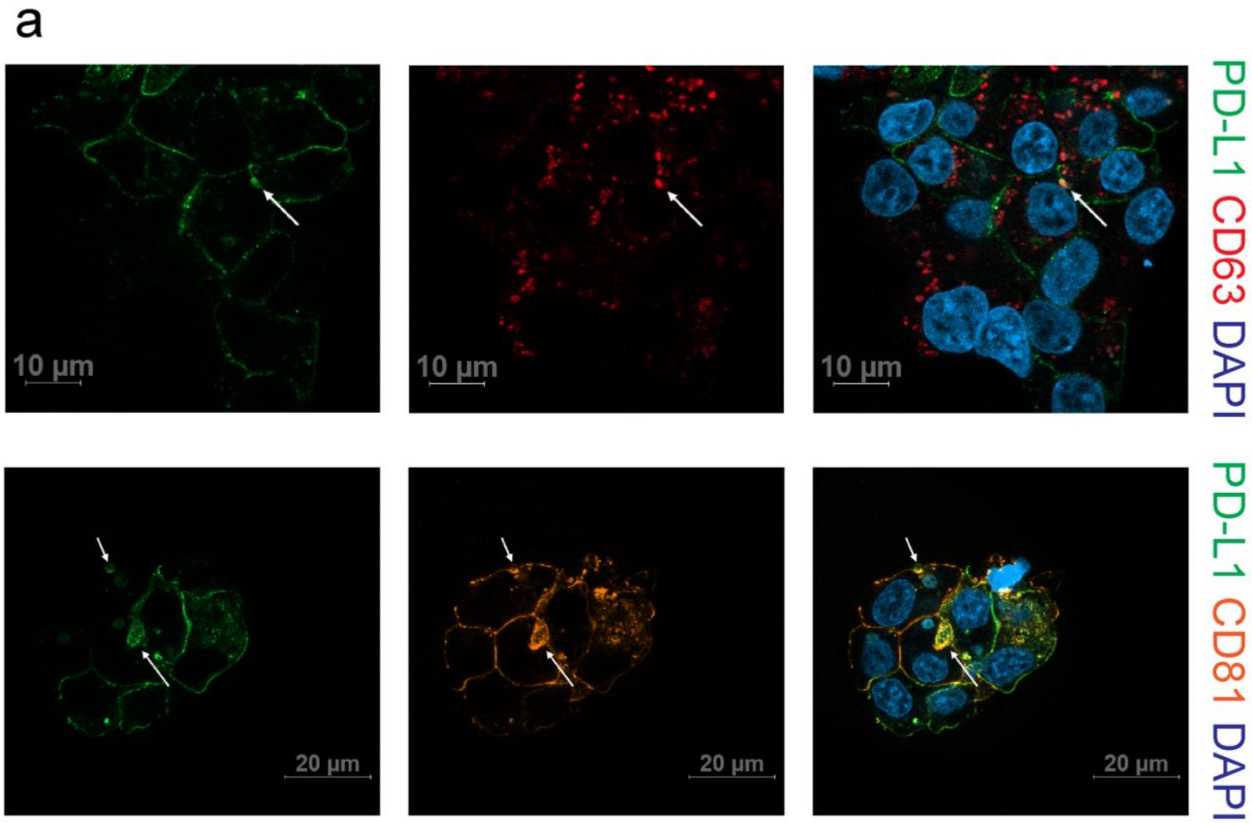


ESM Fig. 1

**PD-L1 colocalizes with beta cell EV-associated proteins in EndoC-βH1 cells. (a)** Confocal microscopy of PD-L1 (green) and tetraspanin-associated surface markers CD63 (red), and CD81 (yellow) in EndoC-βH1 cells.

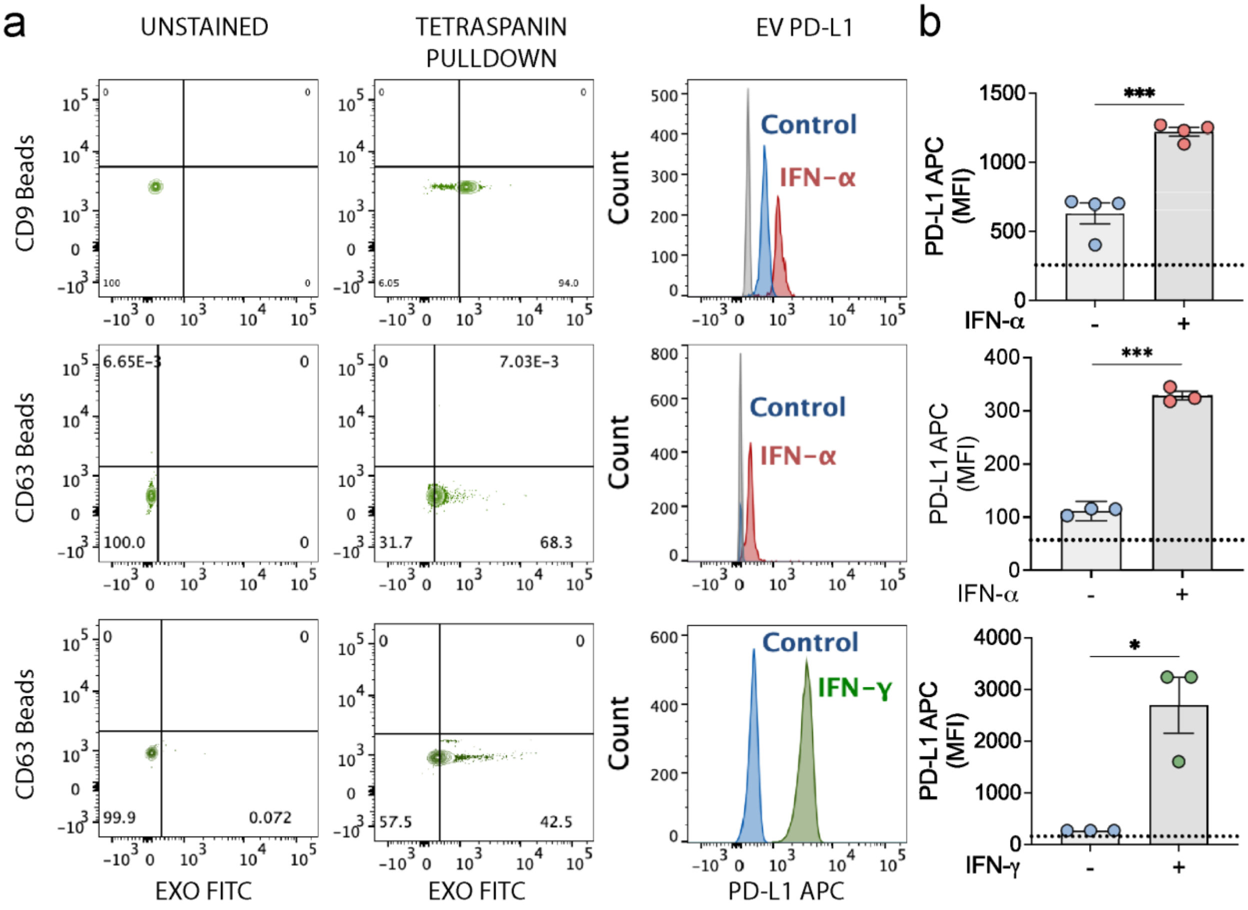


ESM Fig. 2

**PD-L1 protein is present on the beta cell EV surface. (a)** PDL1 EV surface expression as quantified by flow cytometry at baseline or after exposure to IFNα (red) or IFN-ɣ (green). INS-1 EVs +/- 24-h IFN-α were captured with biotin labelled CD9 or CD63 beads and Exo-FITC levels represented as dot plot from unstained or bead pulldown. Histograms represent changes in mean fluorescence intensity of PD-L1 in b (grey), vehicle control (blue), IFN-α treated (red), or IFN-ɣ treated (green). **(b)** The geometric mean of the mean fluorescence intensity (MFI) was quantified at baseline and after exposure to IFNα or IFN-ɣ. Data represented as mean ± SEM. Blue circles indicated vehicle control, red circles indicate IFN-α treated samples, and green circles indicate IFN-ɣ treated samples. Data represented as mean ± SEM; n=3-4; *, p value<0.05; ***, p value<0.001.

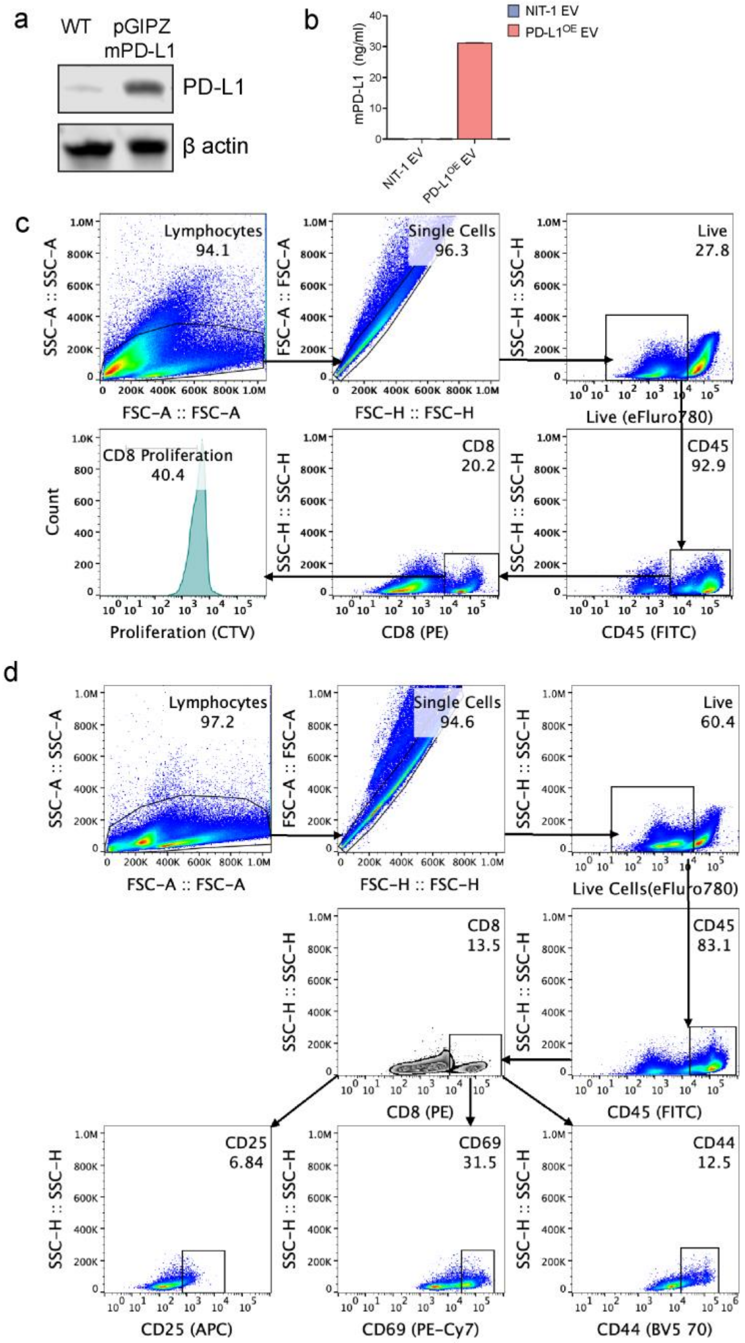


ESM Fig.3

**EV PD-L1 inhibits proliferation, cytokine production and cytotoxicity of murine CD8 T cells *in vitro.*** **(a)** Immunoblot of NIT-1 cells with or without PD-L1 overexpression. β actin is the loading control. **(b)** PD-L1 levels from EVs derived from NIT-1 cells with or without PD-L1 overexpression (PD-L1 OE). **(c-d)** Flow cytometric analysis of CD8 T cells isolated from mouse splenocytes. Gating strategy and representative plots showing **(c)** proliferation assessed by Cell Trace Violet (CTV) and **(d)** activation status of CD8 T cells; determined by CD69, CD25 and CD44 surface expression.

| **Islet preparation** | **1** | **2** | **3** | **4** | **5** | **6** | **7** | **8[a]** |
|---|---|---|---|---|---|---|---|---|
| **MANDATORY INFORMATION** | | | | | | | | |
| Unique identifier | SAMN31815644 | SAMN34411471 | SAMN34997905 | SAMN35085369 | SAMN35334144 | SAMN36458820 | SAMN41162359 | SAMN41474382 |
| Donor age (years) | 32 | 45 | 40 | 32 | 36 | 28 | 54 | 42 |
| Donor sex (M/F) | M | M | M | M | M | M | F | F |
| Donor BMI ($kg/m^2$) | 26.9 | 33.1 | 30.4 | 21.4 | 25.1 | 23.9 | 20.2 | 29.7 |
| Donor $HbA_{1c}$ or other measure of blood glucose control | 4.9 | 5.2 | 5.4 | 5.0 | 5.2 | 4.7 | 5.4 | 5.9 |
| Origin/source of islets[b] | IIDP | IIDP | Alberta Islet Core | Alberta Islet Core | Alberta Islet Core | Alberta Islet Core | Alberta Islet Core | Alberta Islet Core |
| Islet isolation centre | The Schrap-Lacy | University of Wisconsin | | | | | | |

| | | | | | | | | |
|---|---|---|---|---|---|---|---|---|
| | Research Institute | | | | | | | |
| Donor history of diabetes? Please select yes/no from drop down list | No | No | No | No | No | No | No | No |
| **If Yes, complete the next two lines if this information is available** | | | | | | | | |
| Diabetes duration (years) | | | | | | | | |
| Glucose-lowering therapy at time of death[c] | | | | | | | | |
| **RECOMMENDED INFORMATION** | | | | | | | | |
| Donor cause of death | Anoxia | Anoxia | | | | | | |
| Warm ischaemia time (h) | No | No | | | | | | |
| Cold ischaemia time (h) | | | | | | | | |

| | | | | | | | | |
|---|---|---|---|---|---|---|---|---|
| Estimated purity (%) | 90 | 95 | 95 | 90 | 95 | 80 | 75 | 75 |
| Estimated viability (%) | 95 | 95 | | | | | | |
| Total culture time (h)[d] | 78 | 84 | 40 | 39 | 60 | 83 | 16 | 64 |
| Glucose-stimulated insulin secretion or other functional measurement[e] | | | | | | | | |
| Handpicked to purity? Please select yes/no from drop down list | | | | | | | | |
| Additional notes | | | | | | | | |

[a]If you have used more than eight islet preparations, please complete additional forms as necessary
[b]For example, IIDP, ECIT, Alberta IsletCore

[c]Please specify the therapy/therapies
[d]Time of islet culture at the isolation centre, during shipment and at the receiving laboratory
[e]Please specify the test and the results
